# Viral diversity and activity in Arctic tundra soils in wintertime

**DOI:** 10.64898/2026.10.28.762433

**Authors:** Wilhelmiina Juvonen, Marco Forgia, Sirja Viitamäki, Jenni Hultman, Tatiana Demina

## Abstract

Arctic soils store vast amounts of carbon and are warming faster than the rest of the globe. Still, little is known about microbe-mediated feedback responses to the ongoing environmental changes in the warming Arctic. Specifically, viral diversity, functions, and virus-host interactions remain largely unknown in northern soils outside the growing season. Here, we analyzed the diversity and activity of viruses in oroarctic tundra soils during wintertime using metatranscriptomics. Through *de novo* virus prediction as well as targeted searches, we identified 995 viral operational taxonomic units (vOTUs) in total. Most of these vOTUs, 93%, represented RNA viruses, part of which were presumably segments of multipartite genomes. RNA vOTUs were classified mainly within the phyla *Lenarviricota* (259), *Kitrinoviricota* (198), *Pisuviricota* (135), *Ambiviricota* (103), *Negarnaviricota* (79), and *Duplornaviricota* (42). Thirty-seven DNA vOTUs (mostly *Caudoviricetes*) were also detected, suggesting active viral infections. In total, 81% of all vOTUs were assigned with a putative host organism, mostly fungi (515 vOTUs). Viral communities were distinct across vegetation types, and pH and moisture contributed to the differences. Putative novel clades within *Ambiviricota* and *Orpoviricetes* were identified, expanding their known diversity. The study contributes to uncovering diverse RNA viruses and viroid-like elements in Arctic tundra soils and highlights that soil viruses remain active in wintertime.

## Introduction

The Northern circumpolar permafrost region is estimated to contain roughly half of the global carbon stored in soils (Tarnocai et al., 2009). This critical reservoir of carbon is vulnerable to climate change, with the warming across the Arctic occurring up to four times the global rate (Rantanen et al., 2022). Moreover, Northern Fennoscandia has recently experienced a series of record-breaking heatwaves (Rantanen et al., 2025). Soil microbiomes are major drivers of carbon and nitrogen cycling, mediating greenhouse gas emissions, which are expected to increase under warming conditions (Crowther et al., 2019; Maes et al., 2024). Beyond bacterial and archaeal activity, fungi in high-latitude tundra soils have shown an increased capacity to degrade carbon under warming conditions (Cheng et al., 2021). In addition to cellular organisms, viruses are also important players in Arctic soils (Emerson et al., 2018; Trubl et al., 2018), although their input into microbially-mediated carbon transformations remain underexplored, especially outside of the growing season. Despite viral communities have been shown active under simulated winter conditions (Trubl et al., 2021), direct field observations are needed to better understand their ecological significance in wintertime Arctic soils.

Soils harbor a vast diversity of viruses, which can influence the soil microbiome by modulating host population dynamics, thus indirectly affecting biogeochemical cycles (Emerson et al., 2018; Trubl et al., 2018). Through lysing host cells, viruses can restructure microbial communities and release organic matter into the surrounding environment (Carreira et al., 2024). In addition, viruses can influence host metabolism through auxiliary metabolic genes (AMGs) (Jansson & Wu, 2023), for example, by enhancing carbon degradation and thereby affecting ecosystem carbon cycling (Emerson et al., 2018; Wu, Smith, et al., 2022), especially under warming conditions (Ji et al., 2023). Notably, diverse RNA viruses have been detected in Arctic soils and thawing permafrost, where they have been linked to carbon cycling through both virus-encoded metabolic functions and interactions with putative hosts (Pratama et al., 2025; Wu, Bottos, et al., 2022). RNA viral communities are largely associated with fungi, plants, and other eukaryotes and have been shown to vary with environmental conditions and greenhouse gas emissions, suggesting an important role in Arctic ecosystem functioning under climate change (Pratama et al., 2025).

Still, the knowledge of fungal viruses residing in Arctic soils remains fragmentary. Fungal viruses, also called mycoviruses, lack an extracellular phase and are instead transmitted intracellularly via hyphal fusion or spores. Mycoviral infections are often asymptomatic, and the effects on host fungi range from harmful to beneficial (Sutela et al., 2019; Villan Larios et al., 2023). Given this capacity to influence host physiology, mycoviral infection has been suggested to be able to affect fungal decomposition of organic matter (Carreira et al., 2024). Mycoviruses typically have RNA genomes and have proven a rich source of previously unknown RNA virus diversity, with several new lineages described in recent years. Notably, mycoviruses with circular RNA genomes (Forgia et al., 2021; Sutela et al., 2020), established as a novel ribovirian phylum *Ambiviricota* within the kingdom *Orthornavirae*, blur the boundaries between viruses and subviral agents (Forgia et al., 2023; Kuhn et al., 2024). Ambiviral genomes encode an RNA-dependent RNA polymerase (RdRP), a hallmark gene of viruses in the kingdom *Orthornavirae*, but also contain ribozymes typical of viroid-like elements and are thought to replicate via a rolling-circle mechanism (Forgia et al., 2023; Kuhn et al., 2024). Another example of recently discovered mycoviruses is a distinct lineage of ormycoviruses, bisegmented RNA viruses infecting fungi and oomycetes. These viruses are characterized by very diverged RdRPs (Crucitti et al., 2021; Forgia et al., 2022; Olendraite et al., 2023) and are currently classified within the class *Orpoviricetes* in the *Orthornavirae* kingdom (Sabanadzovic et al., 2025).

In this study, we used bulk RNA metatranscriptomics to assess the diversity and activity of viruses in oroarctic tundra soils during wintertime. This approach is particularly suited to detecting fungal viruses, as their intracellular lifestyle makes them difficult to capture through size-based viral enrichment methods (Rodríguez-Ramos et al., 2026). We analyzed viral sequences in 19 metatranscriptomes from soils at Kilpisjärvi, Finland, sampled in winter 2021, representing four vegetation types (deciduous shrub, evergreen shrub, meadow, and fen) (Viitamäki et al., 2025). We hypothesized that wintertime oroarctic soils harbor active and diverse viral communities whose composition is shaped by the same environmental variables that structure bacterial and archaeal communities in these soils (Viitamäki et al., 2025). We also expected to detect a rich fungal virome, including members of recently discovered groups such as *Ambiviricota* and *Orpoviricetes*, whose diversity and ecological significance in Arctic soils remain unexplored. This study expands our understanding of the diversity, ecological roles, and functional potential of viruses in wintertime Arctic soils.

## Materials and Methods

### Sample and metadata collection and sequencing

The dataset comprised samples collected in April 2021 from Kilpisjärvi (Gilbbesjávri in Northern Sámi), in Sápmi, northwestern Finland (69.0479° N, 20.7984° E), covering four vegetation types: evergreen shrubs, deciduous shrubs, meadow, and fen (6, 5, 5, and 3 samples, respectively, Table S1). Sampling strategy, metadata collection, sample processing, and short-read sequencing procedures are detailed in (Viitamäki et al. 2025). Metadata included vegetation type, moisture (gravimetric water content, GWC), pH, soil organic matter (SOM), carbon (C), nitrogen (N), C/N ratio, snow depth and temperature (data available from https://github.com/ArcticMicrobialEcology/kilpisjarvi-winter/blob/main/kilpisjarvi_winter_env_data.csv). Bulk metatranscriptomic data (NCBI BioProject PRJNA1265891; SRA accessions SRR33650596–SRR33650614) were used for assembly and all further analyses.

### Assembly

Raw RNA reads were quality-assessed with FastQC v.0.11.9 (https://www.bioinformatics.babraham.ac.uk/projects/fastqc/) before and after trimming. Trimming was performed using Cutadapt v.4.9 (Martin, 2011), applying the flags ’-m 50 -j 4 – nextseq-trim 20’. *De novo* assembly was carried out with rnaSPAdes v.3.15.5 (Bushmanova et al., 2019) using k-mers 49 and 73, and assembly quality was assessed with rnaQUAST v.2.3.0 (Bushmanova et al., 2016). Reads were mapped back to their respective assemblies using Bowtie2 v.2.5.3 (Langmead & Salzberg, 2012), with SAMtools v.1.21 (Danecek et al., 2021) used for sorting and indexing the resulting alignments.

### Virus prediction and genome annotation

Viral operational taxonomic units (vOTUs) were identified, filtered, clustered, and annotated using the Modular Viromics Pipeline (MVP) v.1.1.4 (Coclet et al., 2024); modules 00-06. For module 01, the databases used were geNomad v.1.7 (Camargo et al., 2024) and CheckV v.1.5 (Nayfach et al., 2021). For module 03, 95% average nucleotide identity and 85% alignment fraction thresholds were used for clustering viral transcripts into vOTUs. In module 05, the flag ’--filtration relaxed’ was applied to retain all vOTUs. Module 06 was run with the flags ’-DRAM’, ’-ADS’, and ’-RdRP’ to generate DRAM-v input and search for anti-defense systems and RdRP profiles, respectively. Only vOTUs of ≥ 1 kilobase (kb) in length were retained for downstream analyses.

High-quality unclassified MVP-derived vOTUs were searched against the IMG/VR v.4 database (Virus DNA DB ver. 4 2022-09-15) using BLASTN with the 1e-5 E-value threshold (Camargo et al., 2023). PHOLD v.0.2.0 (Bouras et al., 2025) was used to additionally annotate prokaryotic MVP-derived DNA vOTUs. The two longest DNA genomes were visualized using clinker (Gilchrist & Chooi, 2021) in CAGECAT v.1.0 (Van Den Belt et al., 2023), with genes colored according to the PHOLD annotations. Putative AMGs were predicted with DRAM-v v.1.5.0 (Shaffer et al., 2020). SegVir v.1 (Tang et al., 2024) was used with the SegVir reference database (DB 2024-05-21) to detect viral segments of MVP-derived RNA vOTUs and vOTUs with an unknown genome type. AMG predictions of interest were further searched with HHPred against the databases PDB_mmCIF70_20_Feb and SCOPe70_2.08, using the MPI Bioinformatics Toolkit (Zimmermann et al., 2018).

In addition to *de novo* viral identification within MVP, targeted searches were used for identifying viruses from the taxa *Ambiviricota* and *Orpoviricetes*. Assembled transcripts were searched against custom RdRP databases for these two viral groups using DIAMOND BLASTX (Buchfink et al., 2015) with an E-value threshold of 1e-10, ’--very-sensitive’ mode, and retaining only the best hit (’-k 1’). For *Ambiviricota*, the database comprised RdRP sequences from (Forgia et al. 2023) (data available at https://github.com/ababaian/serratus/wiki/ambivirus_extended_data). The *Orpoviricetes* database was constructed from all currently accepted *Orpoviricetes* RdRP sequences retrieved from NCBI in March 2026, supplemented with Supergroup24 sequences from (Hou et al. 2024), excluding four narnavirus-like sequences (data available at https://doi.org/10.6084/m9.figshare.26298802.v14). Only hits longer than 1 kb were retained for downstream analyses. The resulting sequences were dereplicated using the CheckV anicalc and aniclust scripts with the thresholds of 95% average nucleotide identity and 85% alignment fraction, following the same approach as for other vOTUs. Hereafter, by vOTUs identified in this study, we refer to all vOTUs recovered using the MVP pipeline as well as through targeted *Ambiviricota* and *Orpoviricetes* searhes. Where analyses were performed exclusively on MVP-derived vOTUs, this is explicitly stated.

### Host prediction

Putative prokaryotic hosts were predicted using iPHoP v.1.4.1 (Roux et al., 2023) for MVP-derived DNA and RNA vOTUs with prokaryotic virus taxonomic predictions, as well as all unknown vOTUs. Host predictions were generated using both the default database (Jun_2025_pub_rw) and a custom database. The custom database was constructed by adding 796 metagenome-assembled genomes (MAGs) previously obtained from Kilpisjärvi soils (Pessi et al., 2022) to the default database. The MAGs were classified with GTDB-Tk v.2.5.2 (Chaumeil et al., 2020) using GTDB release 226 (Parks et al., 2026). The results from the default and custom databases were merged, retaining host taxonomy only to the level at which predictions agreed for the same vOTU. Minimum score thresholds of 75 and 90 were applied for family-and genus-level predictions, respectively.

For MVP-derived RNA vOTUs, host prediction was performed with RNAVirHost v.1.0.5 (G. Chen et al., 2024). vOTUs with order-level taxonomy assigned by MVP were processed directly with the ’predict’ command, while those lacking order-level classification were first run through ’classify_order’ prior to ’predict’. Predictions conflicting between RNAVirHost and iPHoP were discarded.

For *Ambiviricota* and *Orpoviricetes* vOTUs, hosts were assigned manually. All *Ambiviricota* vOTUs were assigned with a fungal host (Chong & Lauber, 2023). For *Orpoviricetes*, the assignments were based on the phylogenetic analysis: vOTUs clustering with the order *Formycovirales* were linked to a fungal host (Sabanadzovic et al., 2025), while the remaining *Orpoviricetes* vOTUs were left unassigned (Petrone et al., 2024).

### Ribosomal RNA removal and read mapping to vOTUs

Ribosomal RNA (rRNA) sequences were identified and removed from trimmed reads using SortMeRNA v.4.3.6 (Kopylova et al., 2012) with the default database (smr_v.4.3_default_db.fasta) applying an E-value threshold of 1e-3. The remaining reads were mapped to all vOTUs using Bowtie2 v.2.5.3 (Langmead & Salzberg, 2012), and the resulting output was sorted and indexed with SAMtools v.1.21 (Danecek et al., 2021). Reads per kilobase million (RPKM) values and covered fraction were computed using CoverM v.0.7.0 (Aroney et al., 2025), applying a minimum read percent identity of 95% and a minimum aligned read percentage of 75%.

### Statistical analysis

Statistical analyses were performed in R v.4.4.2 using the vegan package v.2.6-10 (Oksanen et al., 2025). Principal Coordinates Analysis (PCoA) was conducted using Binary Jaccard distance. In RPKM-based analyses, 10% horizontal coverage threshold was applied. The effects of all environmental variables on viral community structure were assessed by Permutational Multivariate Analysis of Variance (PERMANOVA) with 9999 permutations using Binary Jaccard distances. PERMANOVA was conducted separately for three subsets of the data: all vOTUs, RNA vOTUs, and fungal vOTUs. One sample (271) was excluded from testing the effects of temperature and snow depth through the PERMANOVA analyses due to missing values. Data visualization was performed using the R packages ggplot2 v.3.5.2 (Wickham, 2016), ggalluvial v.0.12.5 (Brunson, 2020), and patchwork v.1.3.1 (https://patchwork.data-imaginist.com).

### Phylogenetic analysis

To assess the phylogeny of *Ambiviricota* and *Orpoviricetes* vOTUs, two separate phylogenetic analyses were conducted. For *Ambiviricota*, sequences with a conserved ABC motif in the RdRP palm domain were selected, alongside ICTV-accepted *Ambiviricota* reference sequences. For *Orpoviricetes*, vOTUs containing ABC motifs in the RdRP palm domain were included together with the complete supergroup24 dataset from (Hou et al., 2024) and ICTV-accepted *Orpoviricetes* reference sequences. Supergroup24 contains viruses that are related to the currently classified members of the order, so it was included in the analyses to have a broader view of the viral diversity within this group. In both cases, sequences were aligned using MAFFT v.7, and maximum likelihood phylogenetic trees were inferred with IQ-TREE v.3.0.1 (Wong et al., 2026) with automatic model selection. Trees were visualized and annotated in iTOL v.7.6 (Letunic & Bork, 2024).

## Results

### Metatranscriptomics revealed taxonomically diverse viral communities

After quality control and trimming, samples retained from 90,242,566 to 522,494,556 reads, with 0.2-3.2% of the reads removed (Table S1). Of the retained reads, 95% were identified as rRNA sequences. Trimmed reads (unsorted) were *de novo* assembled into transcripts, producing from 138,226 to 1,043,424 transcripts per sample. Unsorted reads mapped back to their respective assemblies at the overall alignment rates of 94-98%, which may reflect the high proportion of rRNA reads in the dataset.

In total, 995 vOTUs of ≥ 1 kb in length were recovered using MVP (857) and from targeted searches for *Ambiviricota* (103) and *Orpoviricetes* (35) sequences (Table S2). Out of all vOTUs, 924 and 37 were predicted as RNA and DNA viruses, respectively. The genome type remained unknown for 34 vOTUs. The average length of a vOTU was 2.9 kb, with 2.8, 3.3, and 3.6 kb for RNA, DNA, and unknown vOTUs, respectively. The length of *Ambiviricota* vOTUs averaged to 2.8 kb. Of these, 36 vOTUs fell within the length range of complete *Ambiviricota* genomes (4.3 to 5.2 kb). The average length of *Orpoviricetes* vOTUs was 1.9 kb. The longest vOTU in the dataset was 32.6 kb and represented a DNA virus, while the longest RNA vOTU was 21.6 kb. With the MVP-derived vOTUs, 379 vOTUs were determined by CheckV as complete, high-quality, or medium-quality viral genomes.

Out of the MVP-derived RNA and unknown-genome-type vOTUs, 188 and four were identified as putative segments of multipartite viral genomes, respectively (Table S2). However, segment-to-segment linkage could not be confirmed due to the limited number of samples preventing co-occurrence analysis. In addition, the vOTUs belonging to class *Orpoviricetes* likely represent segmented genomes (note that the sequences recovered here correspond to the RdRP-encoding sequences).

The RNA vOTUs were taxonomically diverse, spanning seven confirmed phyla, with one class-level lineage, *Orpoviricetes*, and one family-level lineage, *Polymycoviridae*, remaining unassigned at the phylum level (Figure 1a). The latter family has been recently placed within the phylum *Pisuviricota* (ICTV, 2026). The phylum *Lenarviricota* accounted for the largest share of vOTUs (259), followed by *Kitrinoviricota* (198), *Pisuviricota* (135), and *Ambiviricota* (103). In addition, RNA vOTUs were assigned to the phyla *Nucleocytoviricota* (79), *Duplornaviricota* (42), unassigned *Orpoviricetes* and *Polymycoviridae* (36), and *Artverviricota* (1). Out of all 924 RNA vOTUs, 71 remained unclassified at the phylum level.

**Figure 1.**
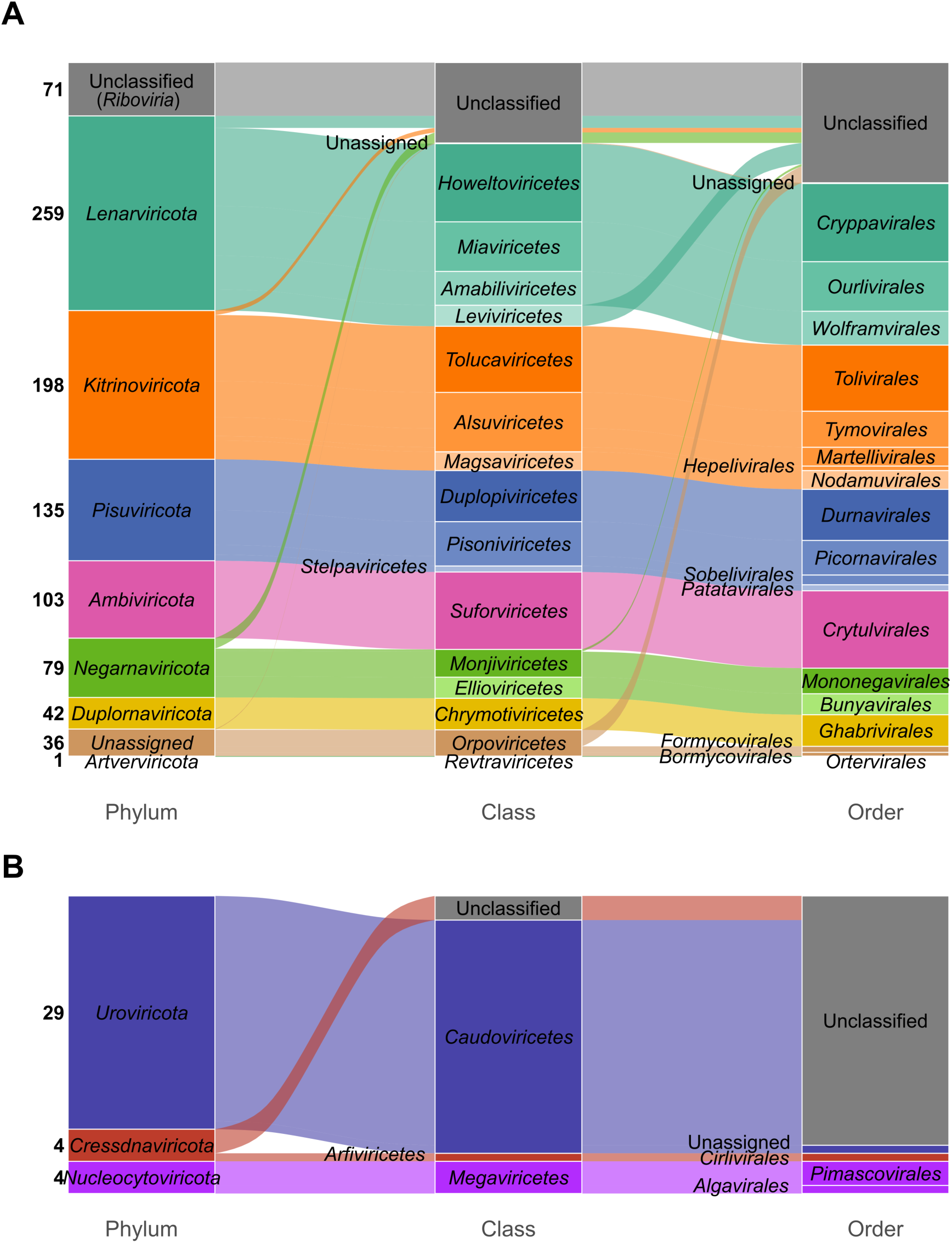
Taxonomic classification of RNA and DNA vOTUs at the phylum, class, and order levels. A) vOTUs with an RNA genome, B) vOTUs with a DNA genome. Note that “Unclassified” refers to vOTUs that could not be classified at this taxonomic rank, while “Unassigned” refers to vOTUs that have lower-level classification, but no higher-level taxon has been defined yet. One order level unassigned *Caudoviricetes* vOTU was classified within the family *Herelleviridae*. One phylum, class, and order-level unassigned RNA vOTU was classified within *Polymycoviridae;* this family has been recently placed within the phylum *Pisuviricota* (ICTV, 2026).

Most of the DNA vOTUs belonged to the phylum *Uroviricota* (29) and were further classified at the class level as *Caudoviricetes* (Figure 1b). One of the *Caudoviricetes* vOTUs was a putative provirus. Two other DNA virus phyla were *Cressdnaviricota* (4) and *Nucleocytoviricota* (4). In total, 34 vOTUs remained unclassified and thus, their genome type could not be inferred from taxonomy. Out of these, four were high-quality and complete, and the PFAM annotation within MVP identified at least one RdRP in each of them, suggesting an RNA virus identity. BLASTN searches against the IMG/VR v.4 database further suggested that three of these unknown high-quality genomes were related to UViGs within *Picornavirales* and one within *Cryppavirales* (Table S3).

In terms of relative abundance, most samples were dominated by vOTUs classified as *Lenarviricota*, *Kitrinoviricota* or *Pisuviricota* (Figure 2a). In three evergreen shrub samples, there was a higher relative abundance of *Pisuviricota* compared to other vegetation types. Meadow samples had the lowest proportions of *Pisuviricota* compared to samples of other vegetation types. *Kitrinoviricota* vOTUs (mainly within the class *Tolucaviricetes*) dominated in samples 115 (deciduous shrub), 577 (evergreen shrub) and 12222 (meadow) (Figure 2b). *Megaviricetes* vOTUs were only found in samples 12221 (fen) and 181 (meadow), while *Arfiviricetes* were only in evergreen shrub samples 37 and 577. Meadow sample 181 had the highest relative abundance of *Caudoviricetes* vOTUs. Unclassified vOTUs could be found from all samples, and especially sample 271 (evergreen shrub) had a large proportion of unknowns. A very low number of vOTUs (13) was detected in the fen sample 12221 overall.

**Figure 2.**
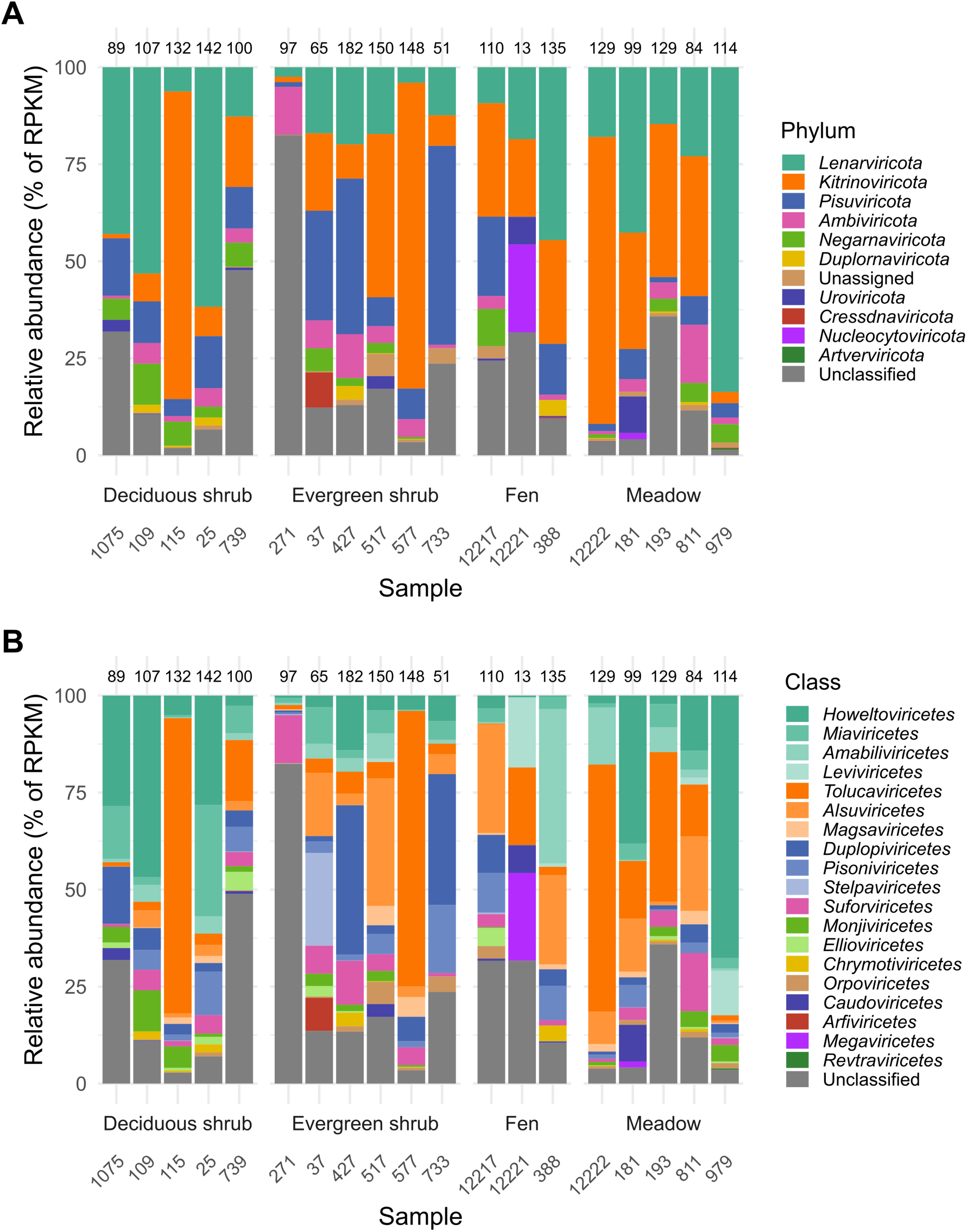
Relative abundance of all vOTUs per sample at the A) phylum and B) class levels. The number of the vOTUs detected per sample is indicated on top of the bars.

### Fungal viruses dominated across the samples

In total, 81% (810/995) of vOTUs were assigned with a putative host by RNAVirHost (676), iPHoP (23), or phylogenetic analysis (111). Over half of all RNA vOTUs were assigned to a putative fungal host (515) (Figure 3a). In addition, invertebrates (179) and Viridiplantae (77) were also common predictions, while Bacteria and Chordata were predicted only for 14 and 8 RNA vOTUs, respectively). Out of all 924 RNA vOTUs, 131 remained with unknown hosts. Four and 13 bacterial hosts were predicted for DNA vOTUs and vOTUs with an unknown genome type, respectively (Figure 3a). However, the majority of DNA vOTUs and vOTUs with an unknown genome type remained without a host prediction (33 and 21, respectively).

**Figure 3.**
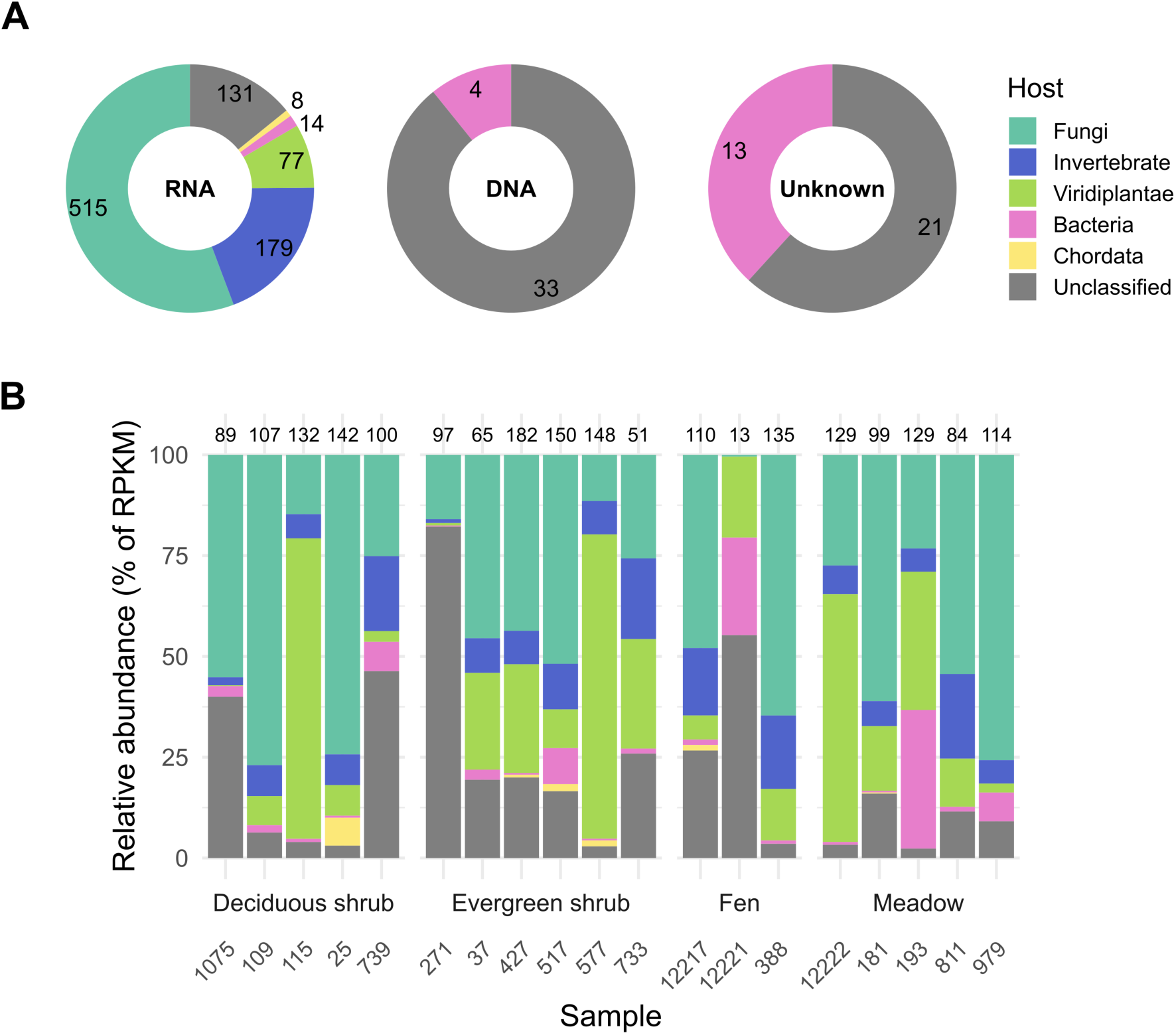
An overview of predicted hosts. A) Number of host predictions for DNA, RNA, and unknown-genome-type vOTUs. The genome type of vOTUs is indicated inside donut plots. B) Relative abundance of vOTUs assigned to different hosts. The number of all vOTUs detected per sample is indicated on top of the bars. The color key is the same for both sections.

Out of all 31 bacterial host predictions, 11, 2, 16 and 2 were domain-, class-, family-, and genus-level predictions, respectively (Table S2). Bacterial host prediction spanned six phyla, with *Bacillota* being the most prevalent (8), followed by *Actinomycetota*, *Bacteroidota* and *Pseudomonadota* (3 each), while *Acidobacteriota* and *Omnitrophota* were each predicted once. Among these predictions, one unclassified RNA vOTU was linked to the family *Bacteroidaceae* (phylum *Bacteroidota*) and four *Orthornavirae* (unclassified further) vOTUs were linked to the families *Acidobacteriaceae* (*Acidobacteriota*), *Bacteroidaceae* (*Bacteroidota*), *Rhodobacteraceae* (*Pseudomonadota*), and *Halomonadaceae* (P*seudomonadota*). Among bacterial hosts predicted for nine *Leviviricetes* vOTUs, only one was further classified within the family *Streptococcaceae* (*Bacillota*) (Table S2). The four DNA vOTUs linked to bacterial hosts were all classified as *Caudoviricetes*, and three of them were predicted to infect hosts within the phylum *Actinomycetota* (families *Mycobacteriaceae*, *Streptosporangiaceae*, and unknown), while one was linked to the family *Enterobacteriaceae* (*Pseudomonadota)*.

We manually assigned all 103 *Ambiviricota* vOTUs to fungi as putative hosts. As for ormycoviruses, viruses within the order *Formycovirales* are known to infect true fungi (Sabanadzovic et al., 2025) and eight vOTUs closely clustering with the representatives of *Formycovirales* were thus assigned to a fungal host (see below for phylogenetic analyses). As *Phormycoviruses* and *Dormycoviruses* have been observed to infect both true fungi and non-fungal hosts such as Oomycetes and *Plasmodium*, no specific host could be assigned to the vOTUs clustering with these taxa.

Relative abundance patterns for vOTUs grouped by predicted hosts varied across samples, though fungal vOTUs were consistently the most prominent (Figure 3b). Fungal vOTUs were present in all samples, and most samples were dominated by vOTUs with fungal, Viridiplantae, or unknown hosts, with no clear differences between samples of different vegetation types. vOTUs linked to bacterial hosts contributed only a minor proportion of relative abundance in most samples, except in the meadow sample 193, where bacterial vOTUs showed the largest proportion (approx. 34 %) across the whole dataset. Invertebrate hosts were observed in all samples except the fen sample 12221 (which showed an unusually low total number of vOTUs compared to the rest of the samples, see above).

### DNA viruses detected across samples suggested that viruses remain active in wintertime soils

DNA viruses were detected across all sample types (Figure S1). Typically, each DNA vOTU was detected in 1-2 samples, but two vOTUs, 517_NODE_1763 and 517_NODE_1809, were present in 4 and 6 samples, respectively. These vOTUs were rather short transcripts (∼2.1 kb), representing *Caudoviricetes*. The median number of DNA vOTUs detected in each sample was 3, but the meadow sample 181 was relatively rich with DNA vOTUs: 19 vOTUs were detected there, including *Caudoviricetes* (17) and *Megaviricetes* (2).

All MVP-derived vOTUs’ predicted ORF products were annotated with geNomad within MVP (Table S3). In total, 1539 functions were predicted, the majority of which either remained unknown (682) (Table S3) or represented RdRPs (660). For DNA vOTUs, 186 geNomad functions were predicted, out of which 129 remained unknown, while 57 ORFs were assigned a specific function. Among the annotated proteins, most were structural components, such as head and packaging proteins (e.g. phage capsid proteins) and tail proteins (e.g. phage tail tube protein, phage tail sheath protein). The remainder were involved in DNA, RNA and nucleotide metabolism (e.g. phage nucleotide-binding proteins, group I intron endonucleases), with small number of other proteins (e.g. phage stabilization protein). In addition to geNomad predictions, *Caudoviricetes* vOTUs were further annotated with PHOLD. In total, 174 ORFs were predicted across the *Caudoviricetes* vOTUs, of which over half had unknown functions (Figure 4a). Of the known functions, the majority fell into three PHROG categories: “DNA, RNA and nucleotide metabolism”, “head and packaging” and “tail” (Figure 4a), reflecting the core replicative and structural components of tailed phages. The remaining annotated ORFs spanned a handful of smaller functional categories.

**Figure 4.**
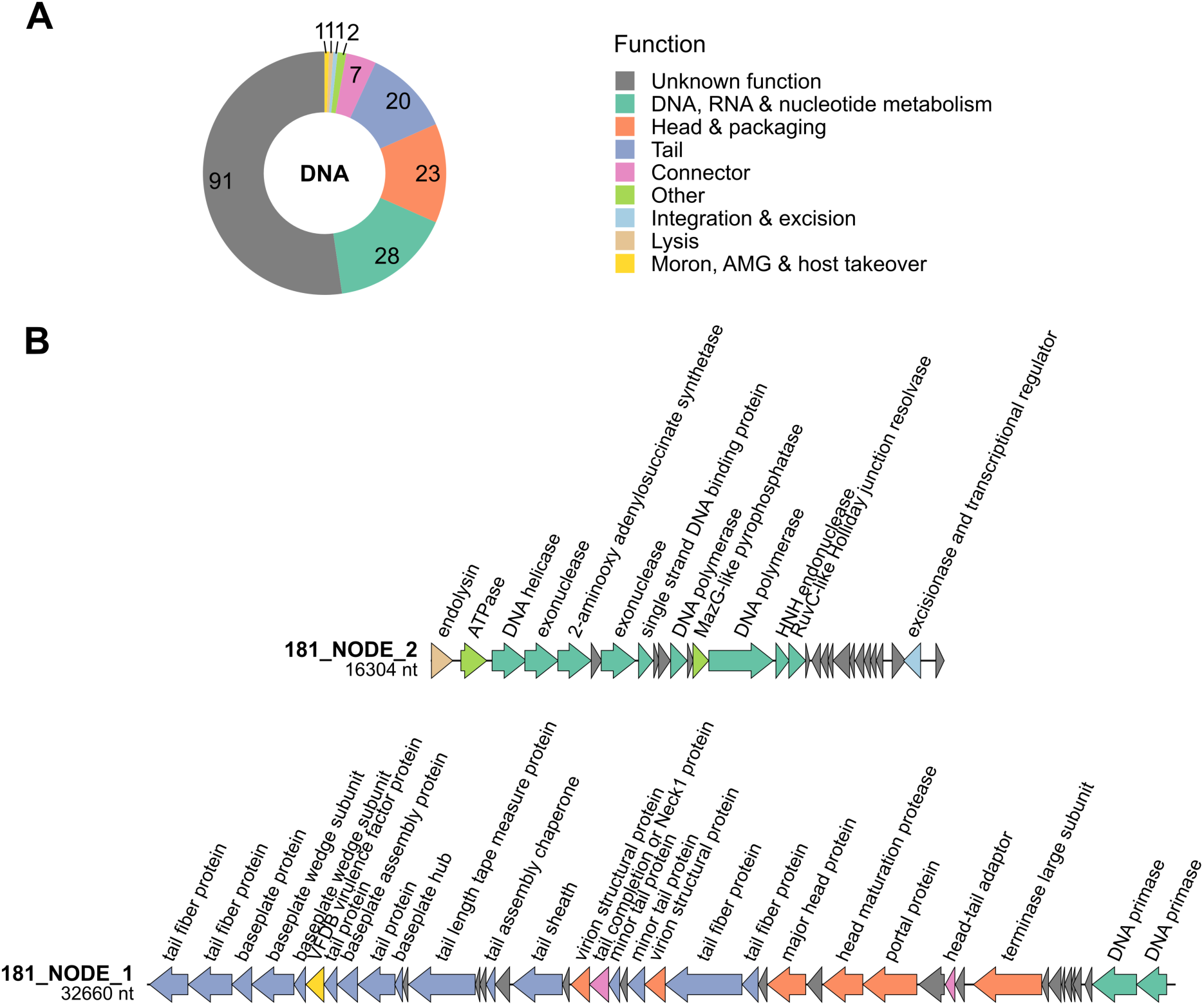
A) The overview of PHOLD functional annotations for all 29 *Caudoviricetes* vOTUs. B) The genomes of the two longest DNA vOTUs (*Caudoviricetes*). ORFs are shown as arrows colored by the functional categories assigned by PHOLD. The color code is the same for both sections. AMG, auxiliary metabolic gene.

Two of the longest *Caudoviricetes* vOTUs, 181_NODE_1 and 181_NODE_2, were 32.6 and 16.3 kb in length, respectively (Figure 4b), with completeness scores of 68.76 and 31.44. These two phages were both present only in the meadow sample 181 and did not share sequence similarity. No hosts were predicted for these two vOTUs. The longest vOTU, 181_NODE_1, encoded various structural genes, including components of the tail (e.g., tail fiber proteins and minor tail proteins), head (e.g., virion structural proteins), as well as connectors (e.g., head-tail adaptor) (Figure 4b). The genome encoded two DNA primases. Notably, one ORF was annotated as a VFDP virulence factor protein and was flagged as a putative AMG. This ORF was annotated as a carbonic anhydrase by PFAM within MVP and additional HHPred searches. The second largest vOTU,181_NODE_2, was a provirus and encoded proteins with functions related to DNA, RNA and nucleotide metabolism, such as DNA helicase, DNA polymerases, exonucleases and a single-strand DNA binding protein. Noteworthy, the 2-aminooxy adenylosuccinate synthase was flagged by DRAM-v as a putative AMG. In addition, the genome encoded an endolysin and an excisionase/transcriptional regulator, consistent with the temperate phage lifestyle capable of switching between lysis and lysogeny.

### Viral communities differed across vegetation types

To explore patterns in viral community composition across samples, PCoA was performed on three subsets of the data: all vOTUs, RNA vOTUs, and fungal vOTUs (Figure 5). In all three subsets, samples clustered by vegetation type, with the two shrub types showing partial overlap. PERMANOVA results showed that vegetation type, along with soil moisture, pH, SOM, C, and C/N ratio, was significantly associated with viral community composition across all three subsets of vOTUs (p<0.05; Figure 5). Vegetation type had the largest effect (R2=0.211-0.228), while soil pH showed the second-largest effect (R2=0.107-0.110). In contrast, N content, temperature, and snow depth were not significant drivers of the community composition in any subset.

**Figure 5.**
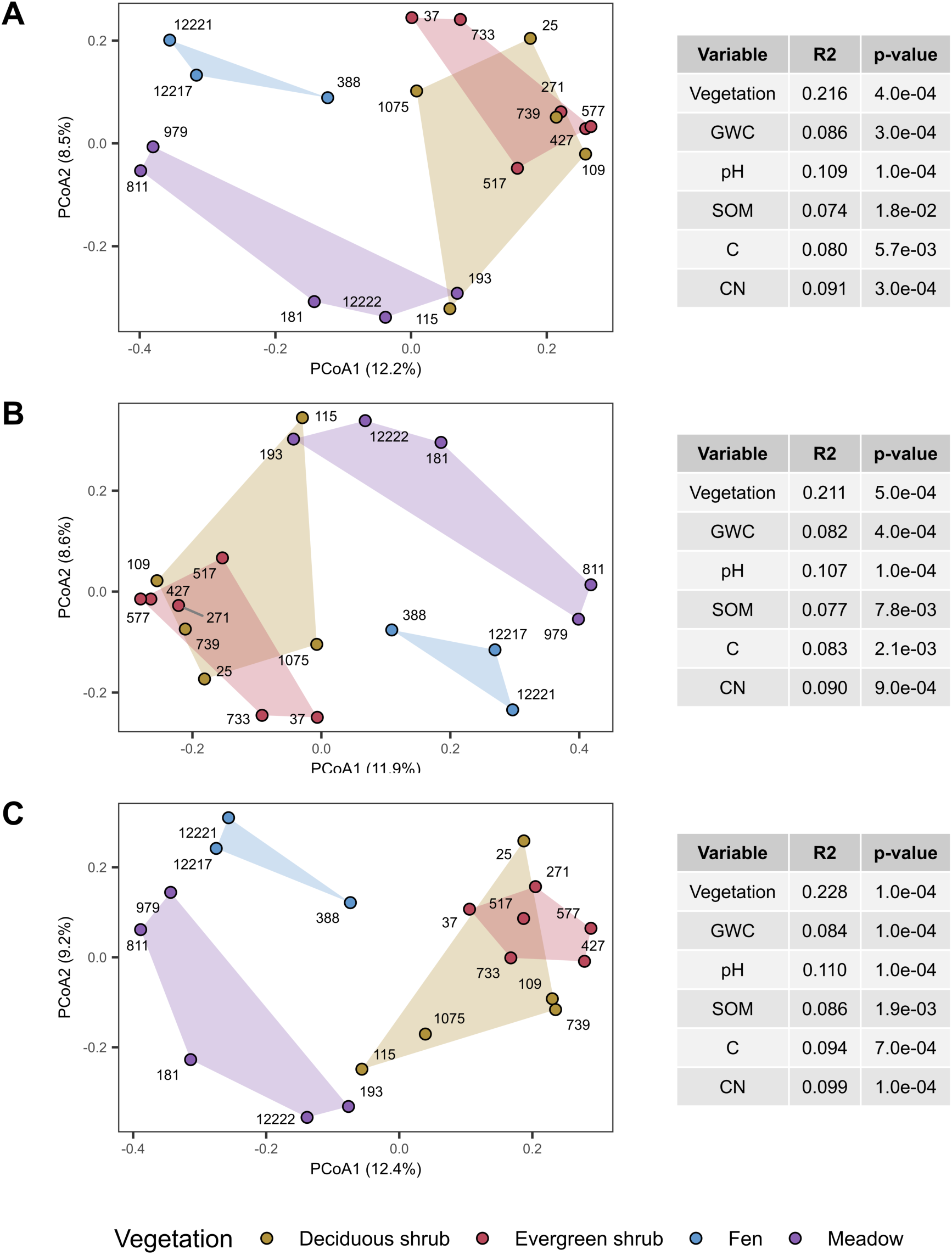
PCoA and significant (p<0.05) PERMANOVA results for A) all vOTUs, B) RNA vOTUs, and C.) fungal vOTUs. A 10% horizontal coverage threshold was used. All parameters were tested separately. GWC, gravimetric water content; SOM, soil organic matter.

### Phylogenetic analyses revealed novel groups of *Ambiviricota* and *Orpoviricetes*

Out of the 103 *Ambiviricota* vOTUs detected, 61 had a conserved ABC motif in the RdRP palm domain and were included in the phylogenetic analysis. The *Ambiviricota* sequences in this study clustered with the representatives of three known genera: *Orthoquambivirus*, *Orthounambivirus* and *Orthodumbivirus* (Figure 6). Viruses falling within the Dumbiviridae family were divided into several clades, indicating that new genera could potentially be established within this family. In addition, some *Ambiviricota* vOTUs recovered in this study formed clades with no known ICTV representatives (Figure 6).

**Figure 6.**
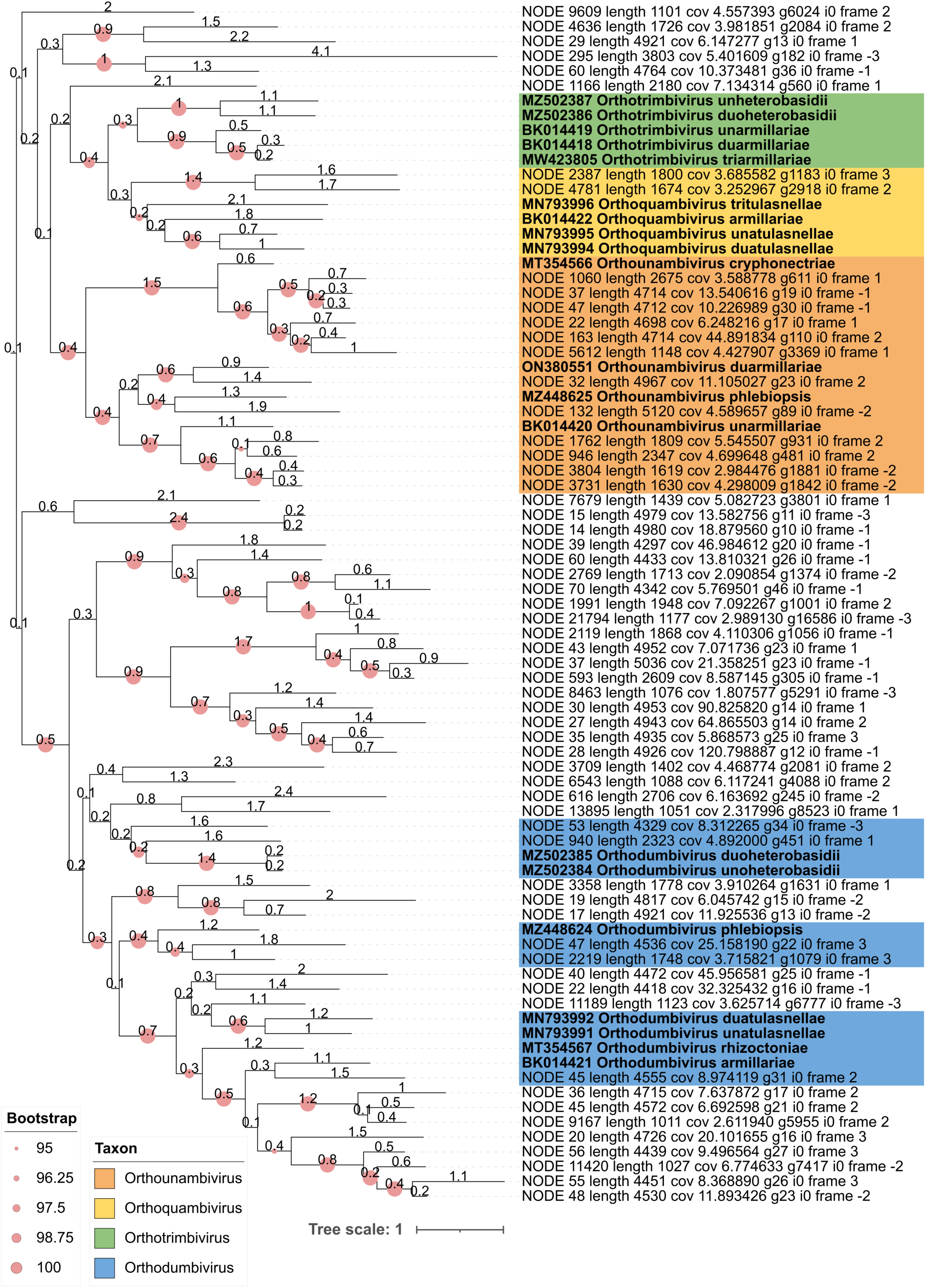
Maximum likelihood phylogenetic tree of the *Ambiviricota* vOTUs identified here as well as ICTV-accepted species (ICTV release 2025; the ICTV-accepted species names are highlighted with bold).

Out of the 35 *Orpoviricetes* vOTUs detected in this study, 21 had a conserved ABC motif in the RdRP palm domain and were included in the phylogenetic analysis. The *Orpoviricetes* vOTUs clustered with the representatives of three known taxa: the order *Formycovirales* as well as the genera *Phormycovirus* and *Dormycovirus* (Figure S2). In addition, several vOTUs clustered together with the Supergroup24 sequences (Hou et al., 2024), with no close ICTV-accepted relatives, suggesting they may represent a previously undescribed taxon.

## Discussion

In this study, we presented 995 vOTUs recovered from Kilpisjärvi oroarctic tundra soil metatranscriptomes. Notably, the analyzed samples were collected from Arctic soils during wintertime (Viitamäki et al., 2025), while most Arctic soil microbiome studies have been focusing on microbial communities during the growing season (Poppeliers et al., 2022). Kilpisjärvi oroarctic tundra soils host diverse bacterial and archaeal communities (Männistö et al., 2024; Pessi et al., 2022), which remain active across seasons (Viitamäki et al., 2022, 2025). These bacterial communities are dominated by *Actinomycetota*, *Pseudomonadota* (*Alphaproteobacteria*), and *Acidobacteriota*, with *Planctomycetota* also abundant, while archaeal communities are largely restricted to *Thaumarchaeota* (Männistö et al., 2024; Pessi et al., 2022; Viitamäki et al., 2022, 2025). As assessed from summertime metagenomes from same sites, DNA viruses are associated with these most abundant groups of bacteria and archaea in Kilpisjärvi soils and carry putative AMGs (Demina et al., 2025).

Among the prokaryotic vOTUs predicted in this study, 31 were linked to bacterial hosts, mostly within the phylum *Bacillota* (8 vOTUs). However, due to a relatively low number of prokaryotic vOTUs detected overall, it is difficult to assess which bacterial phyla they primarily target across samples. Nonetheless, these vOTUs expand the previously described prokaryotic DNA virome of the studied ecosystem (Demina et al., 2025) and include 14 RNA viruses linked to putative bacterial hosts. In addition, linking a few *Caudoviricetes* vOTUs to putative hosts within *Actinomycetota* and *Pseudomonadota* suggest active phage infections that involve abundant bacterial groups in Kilpisjärvi tundra soils across seasons. Furthermore, the longest assembled *Caudoviricetes* vOTU carried a putative AMG, carbonic anhydrase. Carbonic anhydrases are zinc metalloenzymes involved in CO_2_/bicarbonate interconversion and pH regulation and can enhance bacterial growth and host adaptation in some pathogens (Braga et al., 2024). The presence of a carbonic anhydrase AMG in a *Caudoviricetes* genome could suggest a role in host pH control or CO_2_/bicarbonate metabolism, potentially benefiting host fitness and, indirectly, phage replication. However, this remains speculative, as the function of this protein in a viral context is unclear.

Notably, DNA viruses have been observed across all sample types in this study, highlighting their active transcription. To our knowledge, no prior studies report *in situ* DNA virus activity in wintertime tundra soils, though active dsDNA viruses have been detected in Arctic peat soils under simulated winter conditions via stable isotope probing (Trubl et al., 2021). In addition, active dsDNA phages have been reported from high-altitude watershed soils in Colorado during winter (March) under deep snowpack conditions through metatranscriptomics (Coclet et al., 2023). Together, these studies demonstrate that winter conditions can sustain ongoing transcriptional activity of DNA viruses, indicating that soil viral populations remain ecologically relevant during winter months.

The recovered eukaryotic RNA vOTUs spanned a broad taxonomic diversity, with the majority being presumably fungal viruses. As in other soil metatranscriptome studies (Hillary et al., 2022; Pratama et al., 2025) and large-scale survey of >5000 metatranscriptomes across aquatic, terrestrial, and host-associated samples (Neri et al., 2022), most vOTUs fell within the phyla *Lenarviricota*, *Kitrinoviricota* or *Pisuviricota*, with *Negarnaviricota* and *Duplornaviricota* being less abundant. In addition, *Ambiviricota* were also notably abundant in our dataset. Since this phylum, along with the class *Orpoviricetes,* has only recently been established, it is not represented in most of the earlier studies, and its prevalence in soil viromes more broadly remains unclear.

In total, 515 fungal vOTUs reported in this study spanned 6 known phyla and 12 orders, alongside several unclassified and unassigned taxa. Mycoviruses were present in all samples, consistent with previous soil metatranscriptomic studies, where fungi have commonly been predicted as hosts of RNA viruses (Y.-M. Chen et al., 2022; Hillary et al., 2022; Starr et al., 2019). *Lenarviricota* was both the most represented RNA virus phylum in our dataset overall and contained most of the predicted mycoviruses, mainly within the families *Mitoviridae*, *Botourmiaviridae*, and *Narnaviridae* (104, 66 and 45 vOTUs, respectively). *Lenarviricota* viruses have been found abundant in rhizosphere and root litter (Starr et al., 2019), grassland (Wu et al., 2021) and agricultural (Muscatt et al., 2022) soils, watershed sediments (Coclet et al., 2023), and diverse Chinese soils and sediments (Y.-M. Chen et al., 2022). In particular, the prevalence of *Mitoviridae* reinforces findings from grassland soils, where this family was the most frequently detected RNA virus group (later classified within *Narnaviridae* as genus *Mitovirus*) (Starr et al., 2019). It should be noted though that unlike in our dataset, the studies by Coclet et al. (2023) and Muscatt et al. (2022) reported a significant contribution of phages (class *Leviviricetes)* to the phylum *Lenarviricota* in soil viromes. This difference may potentially reflect local environmental conditions across soil habitats.

Among available environmental variables within this study, vegetation type, pH, and moisture were significant drivers of differences in viral communities, including mycoviral ones. In addition, SOM, C, and C/N ratio also contributed to the differences. These observations are consistent with findings from Stordalen Mire, where vegetation type/habitat drove RNA (Pratama et al., 2025) and DNA (Trubl et al., 2025) viral diversity, and pH was the strongest driver of *Orthornavirae* communities (Pratama et al., 2025). Moreover, local habitat has similarly been observed as a key driver of RNA viral community composition across diverse non-Arctic soils (Y.-M. Chen et al., 2022; Hillary et al., 2022; Muscatt et al., 2022; Starr et al., 2019; Wu et al., 2021).

*Ambiviricota* was the second largest phylum of fungal viruses recovered in our dataset, with 103 representatives. This phylum is known to infect both *Ascomycota* and *Basidiomycota*, with *Basidiomycota* reported as the dominant host group (Chong & Lauber, 2023). *Basidiomycota*, *Ascomycota* and *Mucoromycota* dominate the fungal communities of Kilpisjärvi heaths, with *Basidiomycota* being particularly abundant in wintertime (Männistö et al., 2024). These fungal communities are also shaped by habitat and pH (Männistö et al., 2024). Similar patterns observed in shaping host and viral communities, which is also true for bacteria and archaea, whose active wintertime communities are driven by vegetation type and pH (Viitamäki et al., 2025), suggest close ecological coupling between viruses and their hosts in Arctic soils.

Most mycovirus infections are asymptomatic, with no obvious effect on the host phenotype (Sutela et al., 2019). When effects occur, they can, for example, influence the transcription of basic cellular processes or suppress host RNA silencing (Hough et al., 2023; Sutela et al., 2019). Given that mycovirus infections can cause hypovirulence and hypervirulence in their host and thus alter a fungus’s growth rate and competitive fitness (Hough et al., 2023), it is possible that mycovirus infection could modulate the rate of organic matter decomposition in soils. However, with most of the studies on the effects of mycovirus infections being performed on plant-pathogenic fungi, the effects on soil saprotrophic and mycorrhizal fungi remain speculative. At least one *Ambiviricota*, Cryphonectria parasitica ambivirus 1, has been detected to cause hypovirulence in its host (Forgia et al., 2023), whereas the phenotypic effects of *Orpoviricetes* infection remain uncharacterized. As novel viral groups, viroid-like elements, and other RNA replicators continue to be discovered (Crucitti et al., 2021; Forgia et al., 2023; Hou et al., 2024; Sutela et al., 2020), their ecological significance in Arctic soils is still to be uncovered, especially across seasons and in the context of ongoing environmental changes in climate-wise vulnerable ecosystems.

## Data availability

vOTU sequences recovered in this study are available from Figshare: https://doi.org/10.6084/m9.figshare.34296432

## Acknowledgements

We thank CSC – IT Center for Science, Finland, for computational resources.

## Supplementary Figures

**Figure S1.**
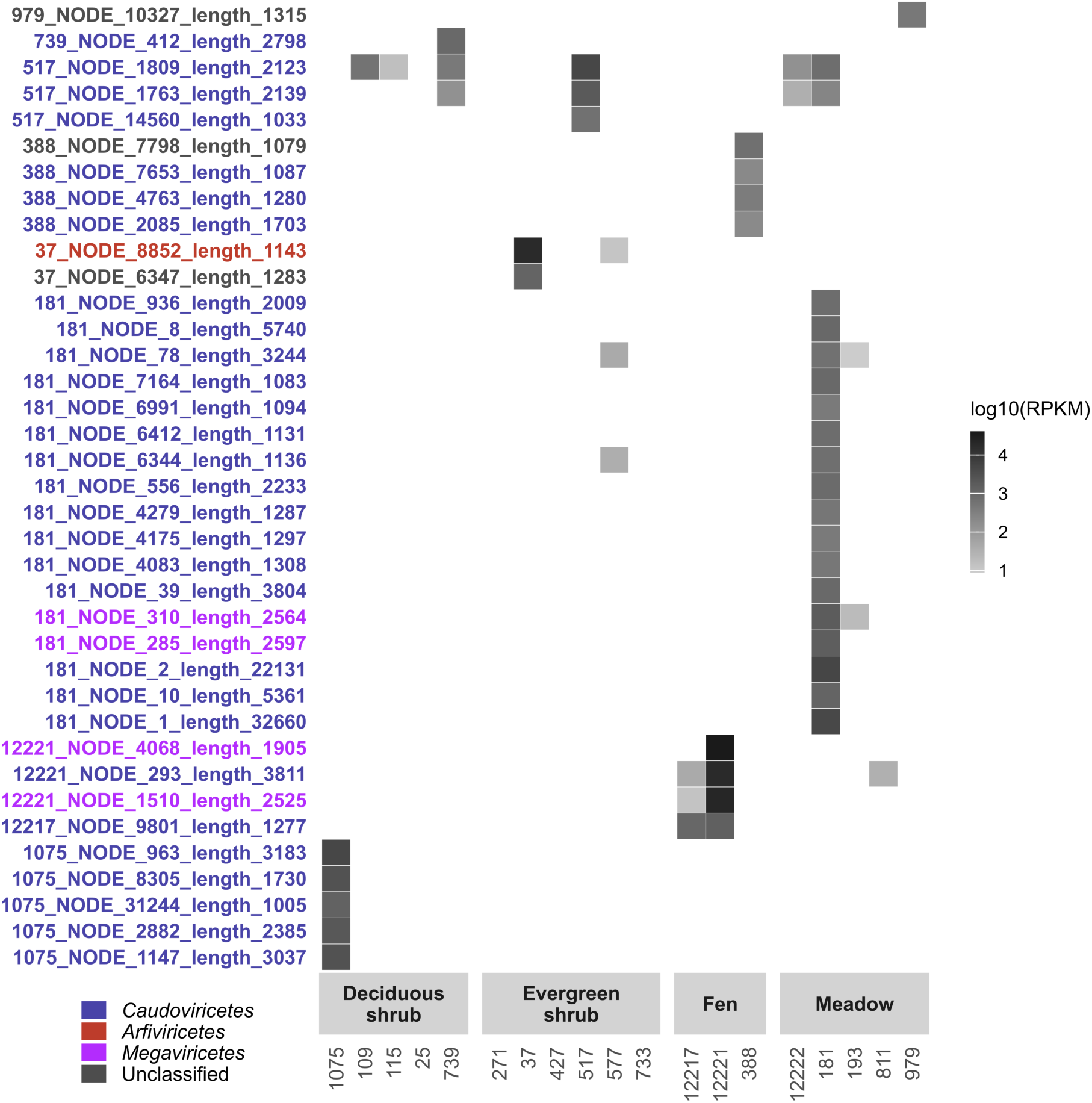
Heatmap of all DNA vOTUs across the samples, 10% horizontal coverage applied.

**Figure S2.**
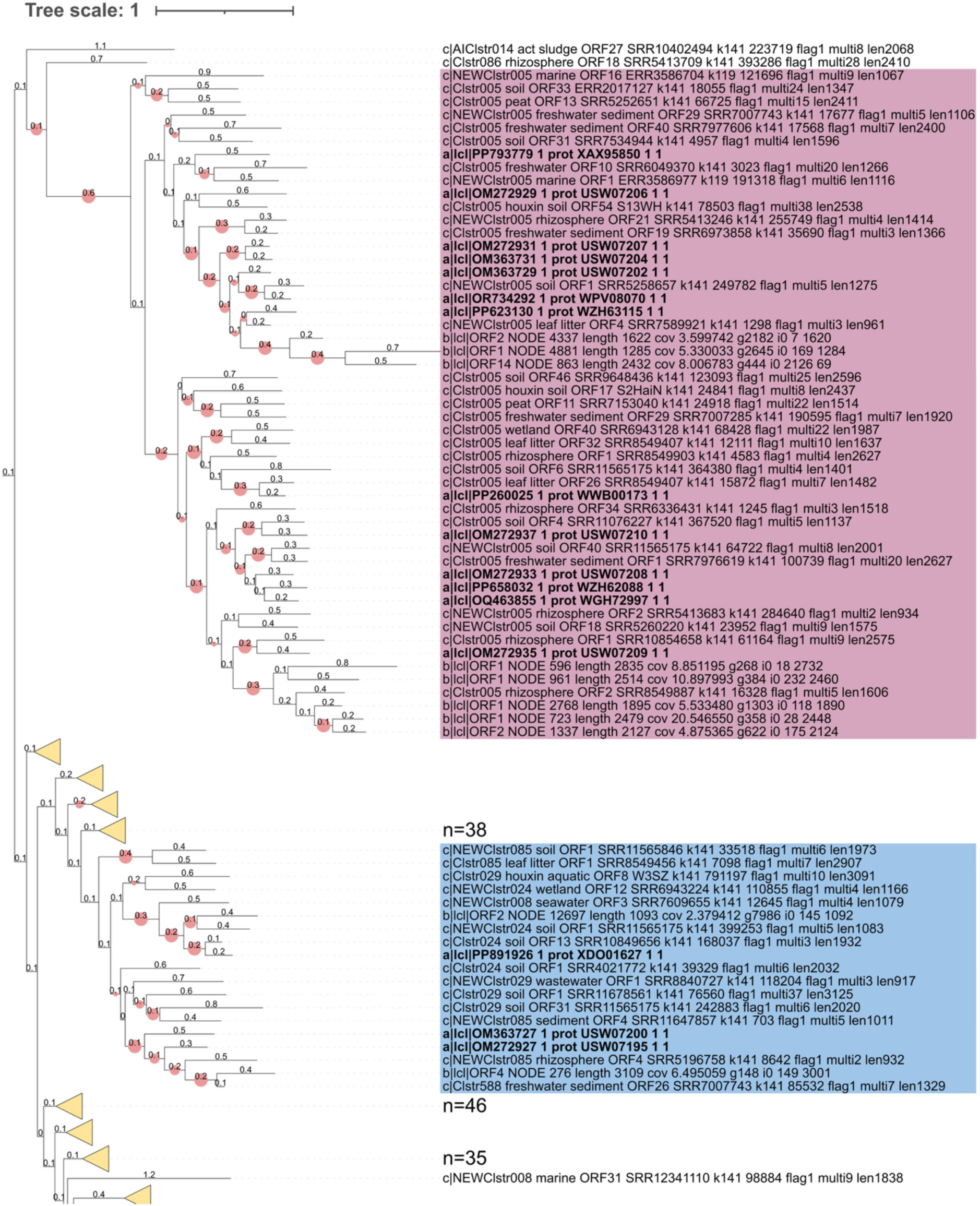
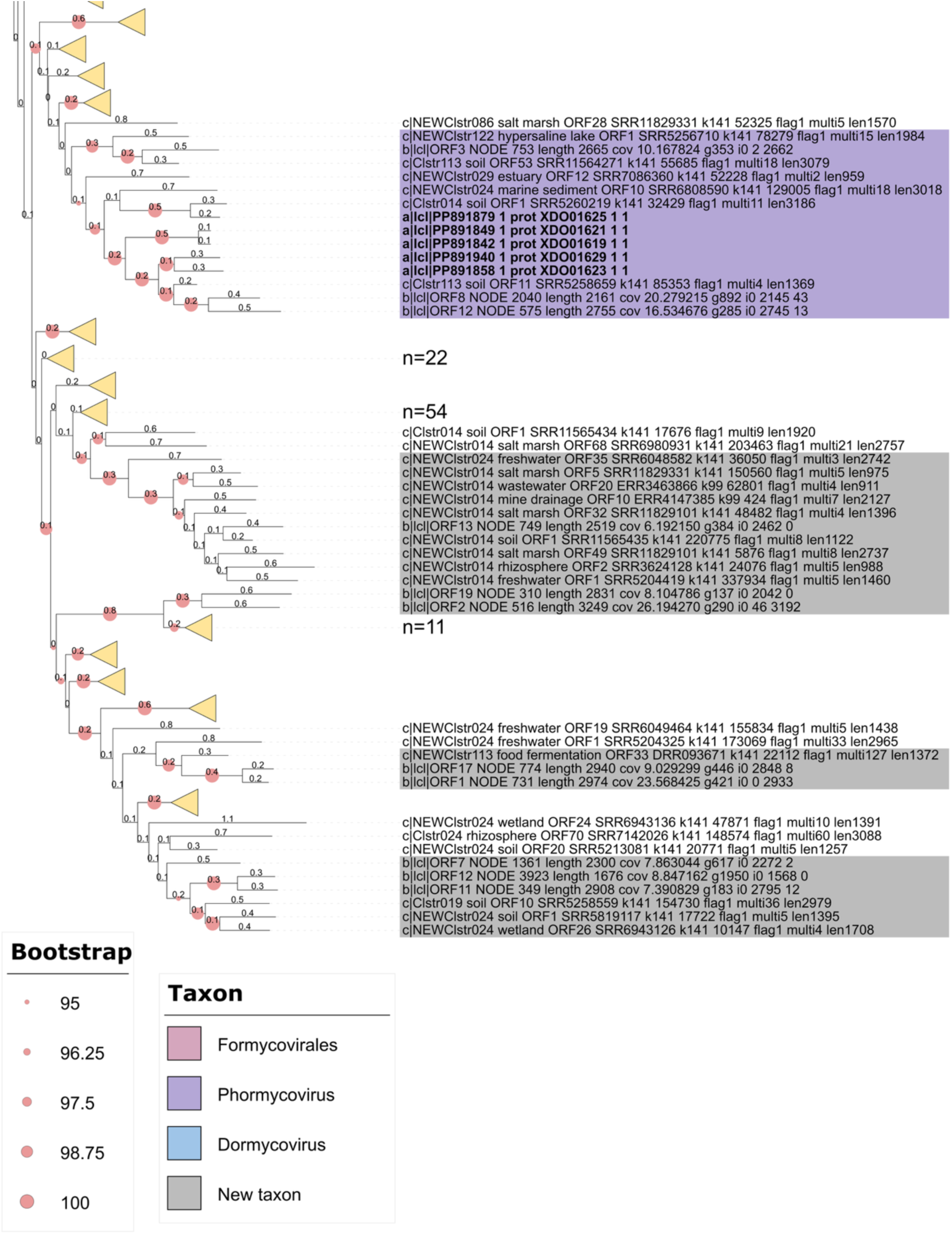
Maximum likelihood phylogenetic tree of the ICTV-accepted *Orpoviricetes* species (labelled with “a” and highlighted with bold), vOTUs from this study (labelled with “b”), and Supergroup 24 (Hou et al., 2024) (labelled with “c”). The numbers of sequences in collapsed clades are indicated for clades that have over 10 sequences.

